# Spatial photosynthesis modelling sets guidelines to constructing a viable single-cell cytoplasm-to-stroma C_4_ cycle

**DOI:** 10.1101/274845

**Authors:** Ivan Jurić, Julian M. Hibberd, Mike Blatt, Nigel J. Burroughs

## Abstract

It has been proposed that introducing C_4_ photosynthesis into C_3_ crops would increase yield. The simplest scheme involves concentrating carbon originating from the cytosol in the chloroplast stroma of mesophyll cells without altering leaf or cell anatomy. Photosynthetic efficiency would then strongly depend on the chloroplast envelope permeability to CO_2_. We examine the performance of this C_4_ cycle with a spatial model of carbon assimilation in C_3_ mesophyll cell geometry, conducting a thorough exploration of parameter space relevant to C_4_ photosynthesis. For envelope permeabilities below 300 µm/s C_4_ photosynthesis has a higher quantum efficiency than C_3_. However, even when envelope permeability is above this threshold, the C_4_ pathway can provide a substantial boost to carbon assimilation with only a moderate decrease in efficiency. Depending on the available light-harvesting capacity of plastids, C_4_ photosynthesis could boost carbon assimilation anywhere from 20% to 100%. Gains are even more prominent under CO_2_ deprivation, and can be achieved in conjunction with lower investment in plastids if chloroplast surface coverage is also altered. A C_4_ pathway operating within individual mesophyll cells of C_3_ plants could hence lead to higher growth rates and better drought resistance in dry, high-sunlight climates.

## Introduction

Various strategies have been proposed to introduce a C_4_ cycle into C_3_ crops with the aim of boosting productivity (Furbank et al., 2013). The evolution of C_4_ photosynthesis is commonly associated with multiple modifications to leaf anatomy, cell biology, and the biochemistry of photosynthesis. Thus, even with rapid advances in our ability to modify plant biology, the endeavour to re-engineer C_4_ photosynthesis is considered challenging. Innovative and versatile theoretical models are therefore needed to guide the design and implementation of carbon concentrating mechanisms in C3 species.

C_4_ photosynthesis is a biochemical carbon-concentrating pump that has evolved over sixty times in higher plants (Sage, 2004). It typically appears in conjunction with so-called Kranz anatomy in which concentric layers of bundle sheath and then mesophyll cells cooperate in the photosynthetic process. Photosynthesis in C_4_ plants is therefore associated with multiple cell walls acting as diffusion barriers to CO_2_ (Sage, 2004). In a small number of species, the C_4_ cycle is contained within individual mesophyll cells (von Caemmerer et al., 2014). It is thought that spatial separation between the primary and the secondary carboxylases (PEPC and RubisCO) in enlarged mesophyll cells phenocopies the diffusion barriers found in two-celled C_4_ plants (von Caemmerer et al., 2014). Introducing a single-cell C_4_ cycle in C_3_ organisms is appealing as the substantial anatomical remodelling of C_4_ leaves and cellular architecture associated with Kranz anatomy could be avoided. However, single-cell C_4_ plants also feature notable modifications to the architecture of mesophyll cells, which facilitate the larger spatial separation (von Caemmerer et al., 2014).

Spatial separation between PEPC and RubisCO aids the C_4_ pump by providing increased diffusive resistance and essentially underpins single-cell C_4_ efficacy (Jurić et al., 2017). However, it is not clear if such cell-scale spatial separation is strictly necessary. To investigate this, we develop a spatial model of a minimal C_4_ pathway operating in an unaltered C_3_ mesophyll cell geometry. The pathway would draw carbon from the cytoplasm and concentrate it within the chloroplast stroma. It would require targeted expression of the pathway enzymes in the cytoplasm and the stroma, a change in the expression of transporters in the chloroplast envelope to transport C_3_ and C_4_ acids, and a C_4_ regulatory mechanism to switch it off when energy/reductant availability is low.

This minimal C_4_ photosynthetic system has previously been discussed by von Caemmerer and Furbank (von Caemmerer and Furbank, 2003; von Caemmerer, 2003) who modelled it within a compartmental paradigm. Their conclusions suggested that although a C_4_ cycle could result in higher CO_2_ assimilation rates, this would come at the expense of a substantially lower energetic efficiency of photosynthesis. However, their analysis assumed a relatively high conductance of the chloroplast envelope, the cell wall, and the plasmalemma (see Discussion). Due to a small spatial separation (∼ 1 *µ*m) between the carboxylase and decarboxylase of the proposed C_4_ pump (which is well below the threshold separation (∼ 10 *µ*m) for cost efficient single-cell C_4_ photosynthesis (Jurić et al., 2017)) the viability of a single-cell based system would be strongly influenced by the permeability of the chloroplast envelope since this determines the CO_2_ leakage current.

CO_2_ permeabilities of biological barriers are uncertain, with estimates of chloroplast envelope permeability ranging over three orders of magnitude, 10^1^-10^4^ *µ*m/s (Evans et al., 2009; Kaldenhoff et al., 2014). We deal with the issue of important input parameters that are poorly characterised by modelling C_4_ photosynthesis for the entire range of reported values. Even so, in order to determine if the proposed C_4_ cycle is viable, we need a criterion to decide whether a particular choice of parameter values is reasonable. One straightforward and simple criterion is that the parameter choice can replicate the known quantum efficiency of regular C_3_ photosynthesis in its native geometry (0.05, or equivalently a photon cost ≈ 20/C (Ehleringer and Pearcy, 1983)). We also quantify the ability of a plastid to absorb and utilise photons for carbon assimilation, by requiring that it does not limit C_3_ photosynthesis under normal conditions. This *light-harvesting capacity* is important because it may constrain assimilation of the proposed C_4_ system and limit attainable yields. We examine the physical parameters affecting C_3_ photosynthetic efficiency in greater detail in a separate publication (under review).

The model we use is based on one previously constructed to study single-cell C_4_ photosynthesis in Bienertia (Jurić et al., 2017). It focuses on the effect of intracellular geometry on the diffusive transport of photosynthetically relevant gasses (O_2_, CO_2_, and its hydrated form HCO^-^3). The diffusion of these species is commonly a limiting factor for both C_3_ and C_4_ photosynthesis. Photochemistry and metabolism (light capture, ATP/NADPH production, Calvin cycle, and photorespiration) are assumed to function optimally. The model is similar in some aspects to the 3D model of C_3_ photosynthesis presented in Tholen and Zhu (2011) but there are notable differences. Most importantly, we include C_4_ biochemistry, but we also explicitly treat oxygen’s kinetics and diffusion, whilst utilising the system’s symmetry to reduce the computational burden, permitting a thorough investigation of the parameter space. By examining how photosynthesis is affected by variation in cell geometry or biochemistry, we can determine when the C_4_ cycle is viable and what alterations are needed to make it a beneficial addition.

## The photosynthesis model

The following is a brief outline of the model. Additional details are provided in the Supplementary Material.

### Geometry

A typical C_3_ mesophyll cell has one large central vacuole that occupies the majority of the cell volume with other organelles located around the cell’s periphery. Chloroplasts in particular, press against the cell membrane in regions adjacent to the intercellular airspace (IAS). Their density is high, with around 50%-70% of the cell surface covered by plastids in a roughly hexagonal lattice arrangement (Figure 1(b)) (Ellis and Leech, 1985; Tholen et al., 2008). Much smaller mitochondria can move freely within the peripheral cytoplasm, which we model as a single homogeneous photorespiring compartment.

**Figure 1:**
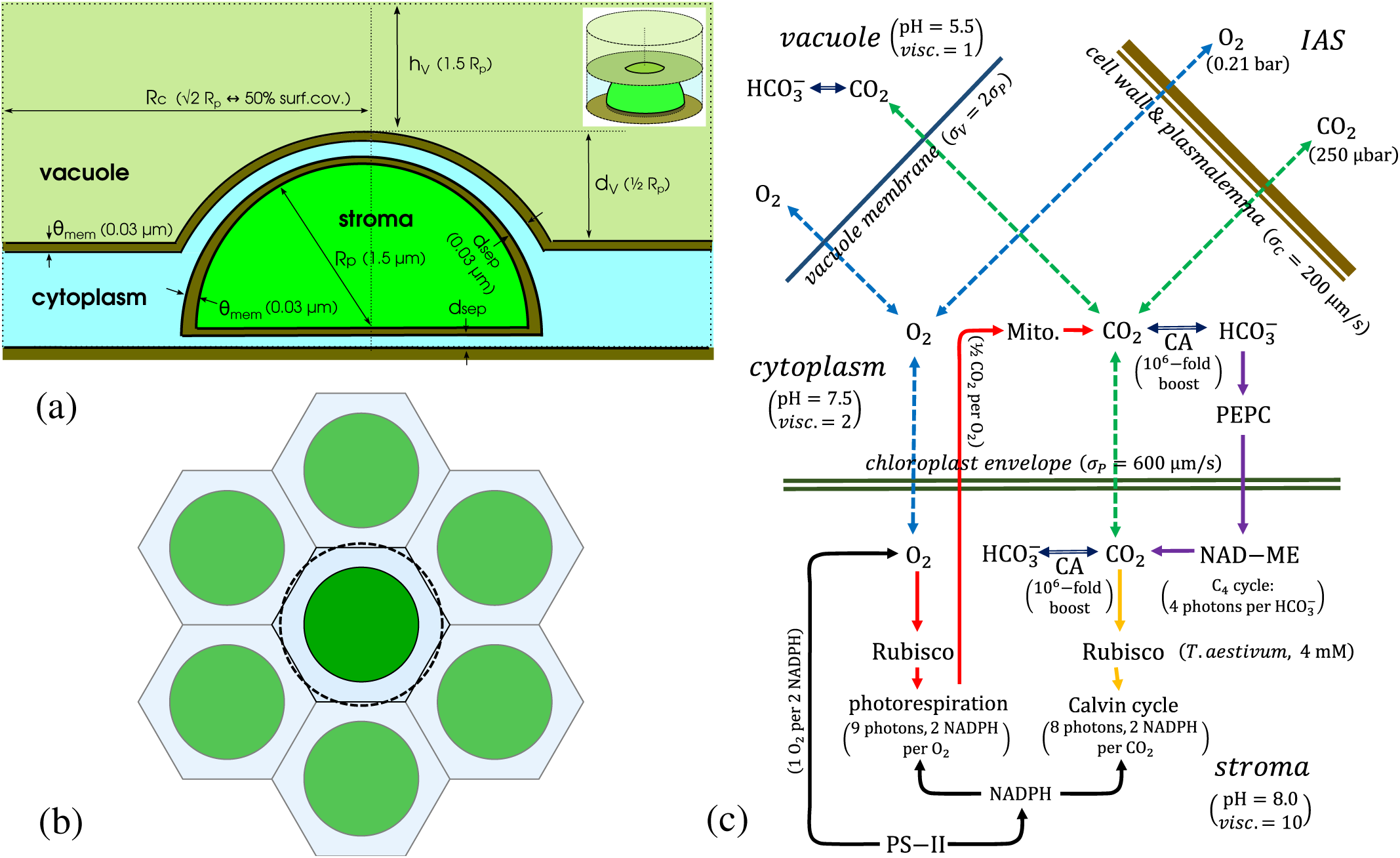
The spatial single-cell C_4_ photosynthesis model. (a): the cross-section of the simulated cylindrical volume (insert) containing a semi- spherically shaped plastid, the peripheral cytoplasm, and a part of the vacuole interior (not to scale). The cylinder radius is determined by the plastid surface coverage. (b): The cylindrical symmetry approximates the ‘personal’ space of an individual plastid in a roughly hexagonal close- packed arrangement of plastids in the areas of mesophyll surface adjacent to IAS. The panel shows such an arrangement at 50% surface coverage ratio. The simulated cylinder is represented by the dashed circle. (c): A schematic representation of the physical processes and chemical pathways modelled. O_2_, CO_2_, and 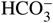 can freely diffuse within individual regions, but O_2_ and CO_2_ can also diffuse through interregional boundaries (dashed green and blue arrows). Depending on the region, the interconversion of CO_2_ and 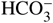 (dark blue arrows) proceeds with or without CA assistance. CO_2_ reacting with RuBP-primed RubisCO drives the Calvin cycle (orange arrows). O_2_ reacting with RuBP-primed RubisCO activates the photorespiratory pathway (red arrows).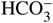 reacting with PEP-primed PEPC is the starting point for the carbon transport through the C_4_ pathway (purple arrows). Oxygen production at PS-II is coupled to the NADPH consumption in the Calvin and photorespiratory cycles (black arrows). Parentheses in (a) and (c) show the default parameter values.

Since both sources and sinks for CO_2_ are located at the cell’s periphery, we expect the vacuole space, especially deeper in the cell interior, to play only a minor role in transport of O_2_ and CO_2_. We therefore focus on a single, typical peripheral plastid and its immediate environment (the spatial region closer to this plastid than to its neighbours), simplifying this approximately hexagonal region as a cylinder (Figure 1(a,b)) that contains one axially-centred semi-spherical plastid. The radius of the cylinder determines the chloroplast surface coverage fraction.

### Transport and biochemistry

We focus on three inorganic species - O_2_, CO_2_, and 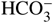- solving for their position-dependent steady-state concentration profiles in order to derive photosynthetic currents. Other metabolites, such as RuBP or PEP, are constrained to the liquid phase and typically do not permeate inter-compartmental boundaries. Their operating pools can in principle be as large as required for optimal photosynthesis, so we assume their levels are not limiting. In contrast, O_2_ and CO_2_ are gasses and readily diffuse within and between cellular compartments, and between the mesophyll interior and outside air. Because of this gaseous exchange, the efficacy of both C_3_ and C_4_ photosynthesis is heavily dependent on their diffusion rates.

Diffusion of the modelled species within particular cellular compartments is quantified by the local viscosities, while diffusion across the inter-compartmental barriers is characterised by barrier permeabilities. Diffusing gasses can enter and exit the simulated region through one of the cylinder ends representing the inner surface of the cell membrane (Figure 1(a)).

Although 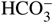 cannot diffuse through intracellular membranes, it too has to be treated explicitly as it strongly couples to the CO_2_ pool in the chloroplast stroma and in the cytoplasm, where we assume CA is present. The CA- assisted interconversion between CO_2_ and 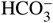is modelled as a boost to the base pH-dependent interconversion rates. This accounts for both the efficiency of CA and its concentration.

The reaction of 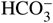with the PEPC-bound PEP in the cytoplasm is the entry point of carbon in the C_4_ cycle. The PEP carboxylation rate determines the rate of CO_2_ release from C_4_-acid decarboxylation in the stroma. The concentration of cytoplasmic PEPC will serve as a measure of C_4_ pathway expression.

RubisCO-bound RuBP in the chloroplast stroma reacts with CO_2_ and with O_2_. The reactions are the key initial steps of the Calvin-Benson cycle and photorespiration. The RuBP oxygenation rate determines the rate of photorespiratory CO_2_ release in the peripheral cytoplasm. The reductant (NADPH) consumption by the Calvin-Benson cycle and photorespiration must be matched by its production via the linear electron transfer chain. This couples O_2_ production on the chloroplast thylakoid with RuBP carboxylation and oxygenation rates.

### Energy input and measures

The assimilation, determined by the throughput of the photorespiratory and Calvin-Benson cycle pathways, is expressed on a cell-surface-area basis. The photon cost of carbon fixation (the number of photons needed per assimilated carbon atom to cover for the costs of the Calvin-Benson cycle, the photorespiration, and the C_4_ cycle) is quantified assuming optimal usage of the linear and cyclic electron transfer chains (Zhu et al., 2008, 2010; Jurić et al., 2017). The total energy consumption is expressed in terms of (photosynthetically active) photons absorbed per stroma volume in unit time, as in Xiao et al. (2016). This consumption cannot exceed the light-harvesting capacity of plastids. As an illustrative reference, a photon consumption rate of 20 mM/s is equivalent to the absorption of 1% of the maximal photosynthetically active solar flux (2 mmol/m^2^s (Björkman and Demmig-Adams, 1995)) that is perpendicularly incident on the parts of the cell surface covered by the plastids (assuming default plastid geometry parameters, Table 1).^1^

**Table 1:** The list of parameters used in the model and in calculation of derived measures. Where not explicitly varied, the parameters are fixed at their default values.

| Parameter | Symbol | Default value | Note |
| --- | --- | --- | --- |
| Plastid radius | $r_P$ | $1.5 \mu\text{m}$ | From Ellis and Leech (1985) |
| Chloroplast surface coverage | $\phi_{P/C}$ | 50% | From Ellis and Leech (1985) |
| Envelope-plasmalemma / envelope-tonoplast membrane separation | $d_{sep}$ | $0.03 \mu\text{m}$ | |
| Envelope and tonoplast membrane thickness | $\theta_{mem}$ | $0.03 \mu\text{m}$ | The membrane thickness is exaggerated to improve numeric convergence. It does not affect the results except through excluded volume. |
| Vacuole drop | $d_V$ | $\frac{1}{2}r_P$ | The depth by which the plastid ‘projects’ into the vacuole space (see Fig. 1(a)). |
| Vacuole height | $h_V$ | $1.5 \times r_P$ | The height (along the central axis) of the simulated part of the vacuole space. |
| RubisCO active site concentration | $c_R$ | 4 mM | Known range is 2 mM-5 mM (von Caemmerer, 2000) |
| PEPC active site concentration | $c_P$ | variable | |
| RubisCO carboxylation catalysis rate | $k_{catRC}$ | $3.8 \text{ s}^{-1}$ | |
| RubisCO oxygenation catalysis rate | $k_{catRO}$ | $0.83 \text{ s}^{-1}$ | |
| RubisCO Michaelis concentration for $\text{CO}_2$ | $K_{RC}$ | $9.7 \mu\text{M}$ | For *T. aestivum* from Cousins et al. (2010) |
| RubisCO Michaelis concentration for $\text{O}_2$ | $K_{RO}$ | $244 \mu\text{M}$ | |
| PEPC carboxylation catalysis rate | $k_{catPbiC}$ | $150 \text{ s}^{-1}$ | |
| PEPC Michaelis concentration for $\text{HCO}_3^-$ | $K_{PbiC}$ | $100 \mu\text{M}$ | For *Z. mays* from Kai et al. (1999) |
| $\text{CO}_2$ pressure in the IAS | $p_{\text{CO}_2}$ | 250 $\mu\text{bar}$ | |
| $\text{O}_2$ pressure in the IAS | $p_{\text{O}_2}$ | 0.21 bar | |
| Henry constant for $\text{CO}_2$ at $20^\circ\text{C}$ | $\nu_C$ | 38.5 mM/bar | From dissolved concentrations at 400 $\mu\text{bar}$ and 210 mbar taken from Carroll et al. (1991) and Murray and Riley (1969). |
| Henry constant for $\text{O}_2$ at $20^\circ\text{C}$ | $\nu_O$ | 1.36 mM/bar | |
| pH in chloroplast stroma |  | 8.0 |  |
| pH in the cytoplasm |  | 7.5 |  |
| pH within the vacuole |  | 5.5 |  |
| $\text{CO}_2 \leftrightarrow \text{HCO}_3^-$ conversion rate boost due to CA | $\eta_{CA}$ | $10^6$ | Saturating, see text. |
| Combined permeability of the cell wall and plasmalemma to $\text{O}_2$ and $\text{CO}_2$ | $\sigma_C$ | 200 $\mu\text{m/s}$ | Ranges in literature from 2 to $5 \cdot 10^3 \mu\text{m/s}$ (Evans et al., 2009; Terashima et al., 2006). |
| Permeability of the chloroplast envelope to $\text{O}_2$ and $\text{CO}_2$ | $\sigma_P$ | 600 $\mu\text{m/s}$ | Ranges in literature from 20 $\mu\text{m/s}$ (Uehlein et al., 2008) to $> 3.6 \text{ cm/s}$ (Missner et al., 2008). |
| Permeability of the tonoplast membrane to $\text{O}_2$ and $\text{CO}_2$ | $\sigma_V$ | $2\sigma_P$ | Assumed to have similar properties to the membranes forming the envelope. |
| Permeability of the chloroplast envelope to $\text{HCO}_3^-$ | | 1 nm/s | |
| Permeability of the tonoplast membrane to $\text{HCO}_3^-$ | | 2 nm/s | Essentially zero. |
| Diffusion constant for $\text{CO}_2$ in water | $D_{C,aq}$ | 1800 $\mu\text{m}^2/\text{s}$ | From Mazarei and Sandall (1980) |
| Diffusion constant for $\text{O}_2$ in water | $D_{O,aq}$ | 1800 $\mu\text{m}^2/\text{s}$ | From Mazarei and Sandall (1980) |
| Diffusion constant for $\text{HCO}_3^-$ in water | $D_{biC,aq}$ | 1100 $\mu\text{m}^2/\text{s}$ | Falkowski and Raven (2013) |
| Cytoplasm viscosity relative to water | $\eta_C$ | 2 | |
| Stroma viscosity relative to water | $\eta_P$ | 10 | As in Tholen and Zhu (2011). |
| Vacuole interior viscosity relative to water | $\eta_V$ | 1 | |
| Plastid light-harvesting capacity | LHC | Varied | Either unlimited, 40 mM/s, or 80 mM/s. |
| Base photon cost of RuBP regeneration | $\varphi_{Calvin}$ | 8 | |
| Base photorespiration photon cost | $\varphi_{phresp}$ | 9 | From Zhu et al. (2010) |
| Base cost of pyruvate-to-PEP conversion | $\varphi_{C4}$ | 4 | |

The limited light availability or light-harvesting capacity is modelled by scaling-down the concentrations of the substrate-primed enzymes involved in photosynthesis (i.e. RuBP-primed RubisCO and PEP-primed PEP-carboxylase), when energy requirements exceed the supply threshold. The adjustment reflects the limited substrate availability caused by energy scarcity.^2^

### The choice of parameters

The default parameters for geometry and biochemistry in Table 1 correspond to *Triticum aestivum* (Ellis and Leech, 1985; Cousins et al., 2010), which we chose as a representative C_3_ crop. Not all parameters are well known however, and some reflect environmental conditions. We therefore examine how the variation in potentially important physical parameters would affect the efficacy of the proposed C_4_ pathway. This is executed by independently varying the selected parameter and the activity of the C_4_ pump (i.e. the cytoplasmic PEPC level). Since the plastid envelope is expected to have a major influence on the efficiency of the pathway, we focus on varying its permeability, while keeping the combined permeability of the cell wall and plasmalemma at σ_*C*_ = 200 *µ*m/s (representing the mid-range of experimental estimates provided by Terashima et al. (2006) and Evans et al. (2009)). The permeability of the vacuole membrane always defaults to twice the envelope permeability (the latter being a double membrane). The actual effect of independently varying the vacuole membrane permeability will be shown to be negligible. The concentration of RubisCO active sites in the stroma is kept at 4 mM. This represents the concentration of activated and RuBP- primed RubisCO, and is roughly in the middle of the known range of RubisCO active site concentration (2-5 mM; von Caemmerer (2000)).

When evaluating the relative efficiency of the C_4_ cycle, we use C_3_ photosynthesis *with the same amount* of CA in the cytoplasm as the baseline for comparison. Since cytoplasmic CA already improves photosynthesis slightly (see Results), using the ‘enhanced’ C_3_ photosynthesis as a baseline benchmark will produce more conservative estimates of gains when introducing a C_4_ cycle.

## Results

We first examine photosynthesis without any imposed light-harvesting cap, only observing at which PEPC concentration a particular energy consumption threshold is reached. Later we examine what happens when an actual light-utilisation cap is imposed.

### The impact of cytoplasmic and stromal CA on photosynthesis

There needs to be sufficient CA in the cytoplasm to allow for a fast conversion of CO_2_ into bicarbonate, which is a substrate for PEPC. CA is known to be present in the chloroplast stroma in C_3_ plants (Tiwari et al., 2005; Tsuzuki et al., 1985). There is also some evidence of cytoplasmic CA expression (Tiwari et al., 2005; Fabre et al., 2007), although the level of its activity and its effect on photosynthesis remains unknown. It has been conjectured that the stromal CA’s purpose is to boost CO_2_ diffusion within the plastid, or to facilitate CO_2_ transfer through the envelope by generating a larger CO_2_ gradient across this diffusion barrier (Badger, 2003). Previous modelling has shown a minor positive impact on the assimilation rate attributable to stromal CA (Tholen and Zhu, 2011). Our results support these findings, showing an increase to C_3_ photosynthetic efficiency and assimilation rate at CA conversion efficiencies (*η*_*CA*_) above 10^3^ (Figure 2). The gain reaches 10% at *η*_*CA*_ = 10^6^ and begins to saturate at larger *η*_*CA*_. Interestingly, the effect is essentially independent of the envelope permeability value, as long as we are not close to the compensation

**Figure 2:**
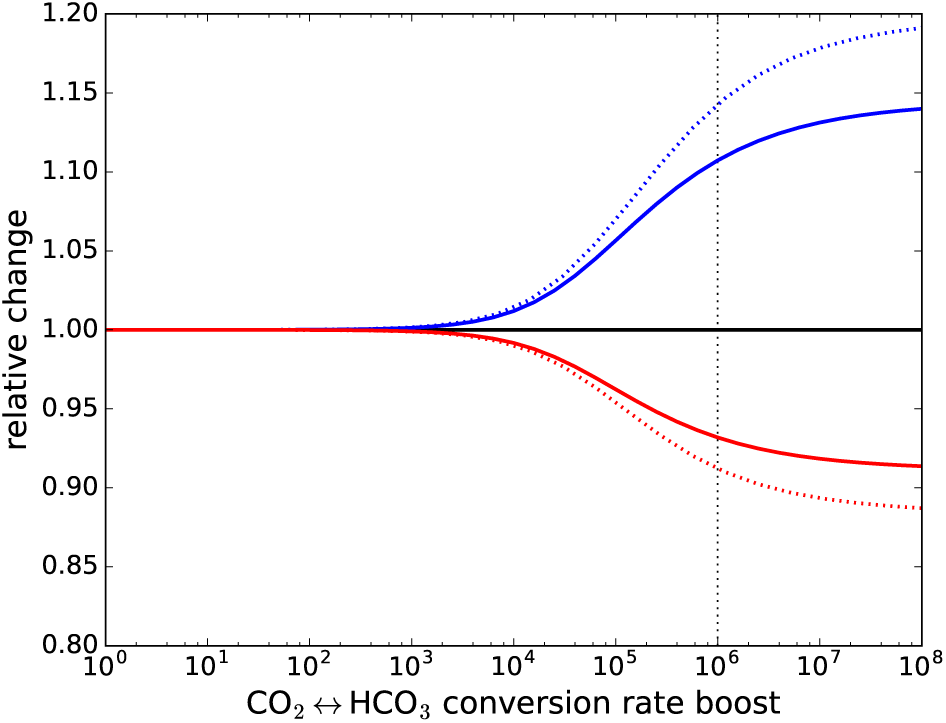
Cytoplasmic CA and C_3_ photosynthesis. The assimilation rate gain (blue) and the photon cost reduction (red) as functions of CA effectiveness (*η*_*CA*_) for σ_*P*_ = 600 *µ*m/s. Solid lines are for the case of CA present only in the plastid stroma; dashed lines are for the case of CA present both in the stroma and in the cytoplasm. The vertical dotted line marks the boost factor used as default in other figures.

point (Supplementary Figure 1). The results are similar when CA is expressed both in the chloroplast stroma and in the cytoplasm, but with a somewhat larger increase in C_3_ efficiency and assimilation (∼ 14% at *η*_*CA*_ = 10^6^, Figure 2).

With a C_4_ cycle expressed, changing the efficacy of the cytoplasmic CA^3^ can greatly affect photosynthesis (Supplementary Figure 2). Cytoplasmic CA activity acts as one of the bottlenecks to the pump throughput; for *η*_*CA*_ < 10^4^ the pump is effectively non-operational and varying the PEPC level produces no noticeable change in the photon cost or the assimilation rate. For *η*_*CA*_ beyond 10^6^, CA ceases to be a limiting factor at PEPC concentrations below 1 mM. Hence, we select *η*_*CA*_ = 10^6^ as a default value both for cytoplasm and stroma. Depending on how effective the CA strain is, an *η*_*CA*_ of 10^6^ would correspond to a CA active site concentration of 0.2 mM (spinach CA; Pocker and Ng(1973)) or ∼ 1 mM (pea; Johansson and Forsman (1994)).

### The impact of the gas permeability of the plastid envelope

We next investigate how the photon cost and assimilation rate depend on the envelope permeability and the PEPC concentration (i.e. the pump activity), Figure 3. Both photon cost and assimilation rate begin to change notably when PEPC concentration reaches 10^−2^ - 10^−1^ mM. By 1 mM, the efficacy measures saturate, as the pump reaches full activity. Taking into account the volume of the plastid and the surrounding cytoplasm, the PEPC concentration range of 10^−2^ - 10^−1^ mM corresponds to a PEPC-to-RubisCO carboxylation capacity ratio between 0.1 and 1, while saturation occurs at ratios close to 10. By comparison, the PEPC/RubisCO activity ratio in C_4_ plants is between 2 and 6.5 (von Caemmerer et al., 2014). The reason behind the saturation in the photosynthetic activity at high PEPC concentrations is a limitation in the carbon supply - either the CA-assisted 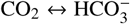conversion rate becomes insufficient or the diffusion of CO_2_ from IAS through the cell wall reaches its limit.^4^ Realistically however, we can expect that energy expenditure will limit photosynthesis before that, as we demonstrate later.

**Figure 3:**
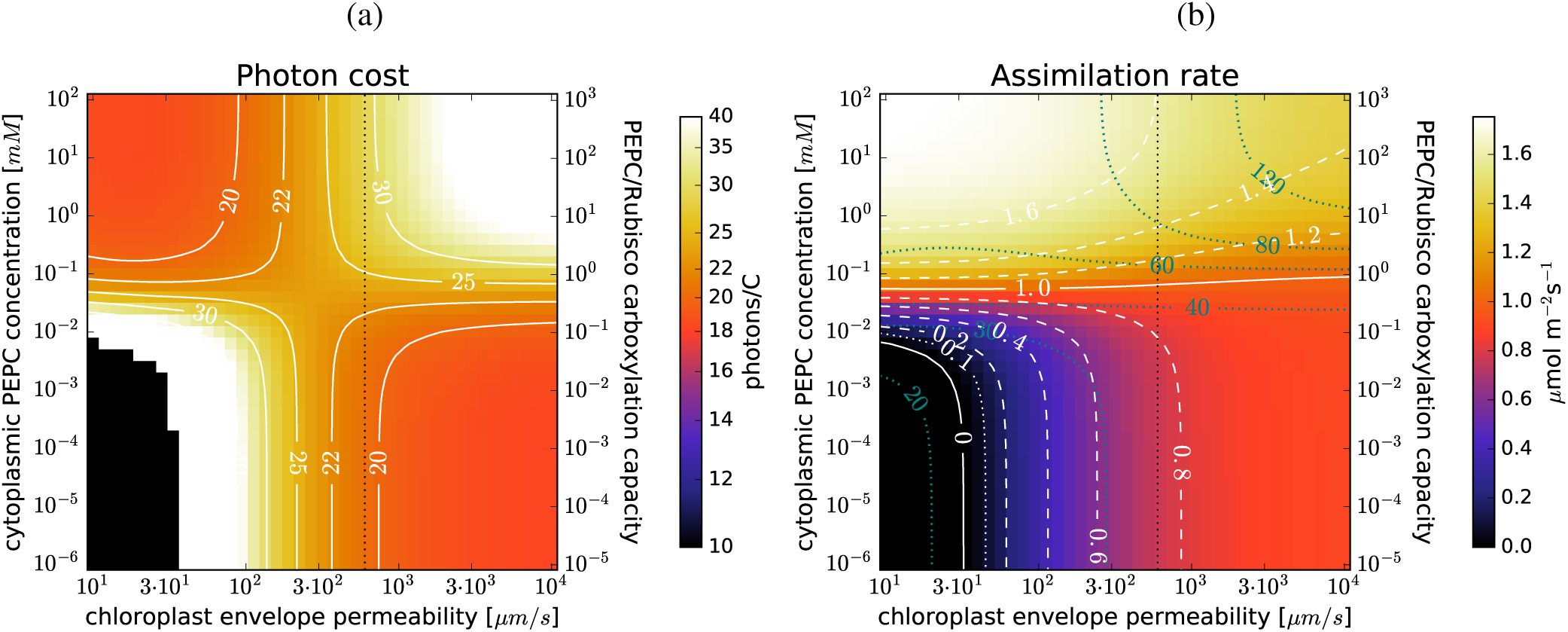
Envelope permeability and C_4_ photosynthesis: The photon cost (a) and the net assimilation rate (b) as functions of the envelope per- meability and PEPC concentration in the cytoplasm for the default parameter choice (Table 1). The purple lines in (b) mark the light-utilisation thresholds (in mM/s). In the black regions the photon cost and the assimilation rate are negative. The vertical dotted line marks the envelope permeability used as default in other figures.

Of particular interest is how the photon cost changes with the pump activity level at different permeability values. At envelope permeabilities (σ_*P*_) below a threshold value of ∼ 300 *µ*m/s, the photon cost decreases when the pump is operational. Above the threshold permeability the opposite happens. At low envelope permeabilities the C_4_ pathway would therefore be cost-efficient compared to C_3_ photosynthesis. However, this regime is unlikely to occur in reality. To confirm this we can look at the photon cost at negligible PEPC concentration (that is, during C_3_ photosynthesis). With the cell boundary permeability fixed at 200 *µ*m/s, the photon cost of C_3_ photosynthesis reaches 20/C for envelope permeability σ_*P*_ ≈ 600 *µ*m/s (Figure 3), i.e. the actual permeability of the chloroplast envelope is higher than the 300 *µ*m/s threshold and the photon cost of C_4_ photosynthesis is higher than of C_3_ photosynthesis. We note however that the threshold permeability value, where the C_4_ pathway breaks-even, changes with CO_2_ pressure in the internal airspaces, moving to higher values as the pressure decreases (see Supplementary Figure 3). Consequently, even for realistic envelope permeability values (i.e. several hundred *µ*m/s) the proposed pathway *can* become a cost-efficient strategy under conditions of CO_2_ deprivation (IAS CO_2_ pressure *p*_*CO*_2 < 150 µbar, such as may occur during prolonged stomata closure).

Although the C_4_ cycle may not be cost-effective in terms of quantum efficiency, it always increases the assimilation rate at sufficiently high PEPC activities. The gain can be substantial - up to several-fold at high PEPC concentrations assuming photosynthesis is not limited by light (Figure 3(b)). The light-harvesting capacity of a plastid can be established by looking at the energy consumption of C_3_ photosynthesis when the photon cost is not larger than 20/C (σ_*P*_ 600 *µ*m/s). We can see from Figure 3(b) that the light-harvesting capacity should be larger than 30 mM/s of photosynthetically active photons per stromal volume. We take 40 mM/s as a rough estimate of the actual, or at least easily achievable light-harvesting capacity of an average plastid. As we demonstrate later, significant assimilation gains are feasible even at this limited capacity.

### The impact of other variables

The impact of the vacuole membrane permeability on the C_4_ cycle efficiency is minimal (Supplementary Figure 4), although changing the thickness of the peripheral cytoplasmic layer does change the cytoplasmic PEPC concentration at which a particular efficiency or gain level is achieved (Supplementary Figure 4(c)). It appears that what really matters is the total amount of PEPC in the cytoplasm compared with RubisCO in the plastids (Supplementary Figure 4(d)).

**Figure 4:**
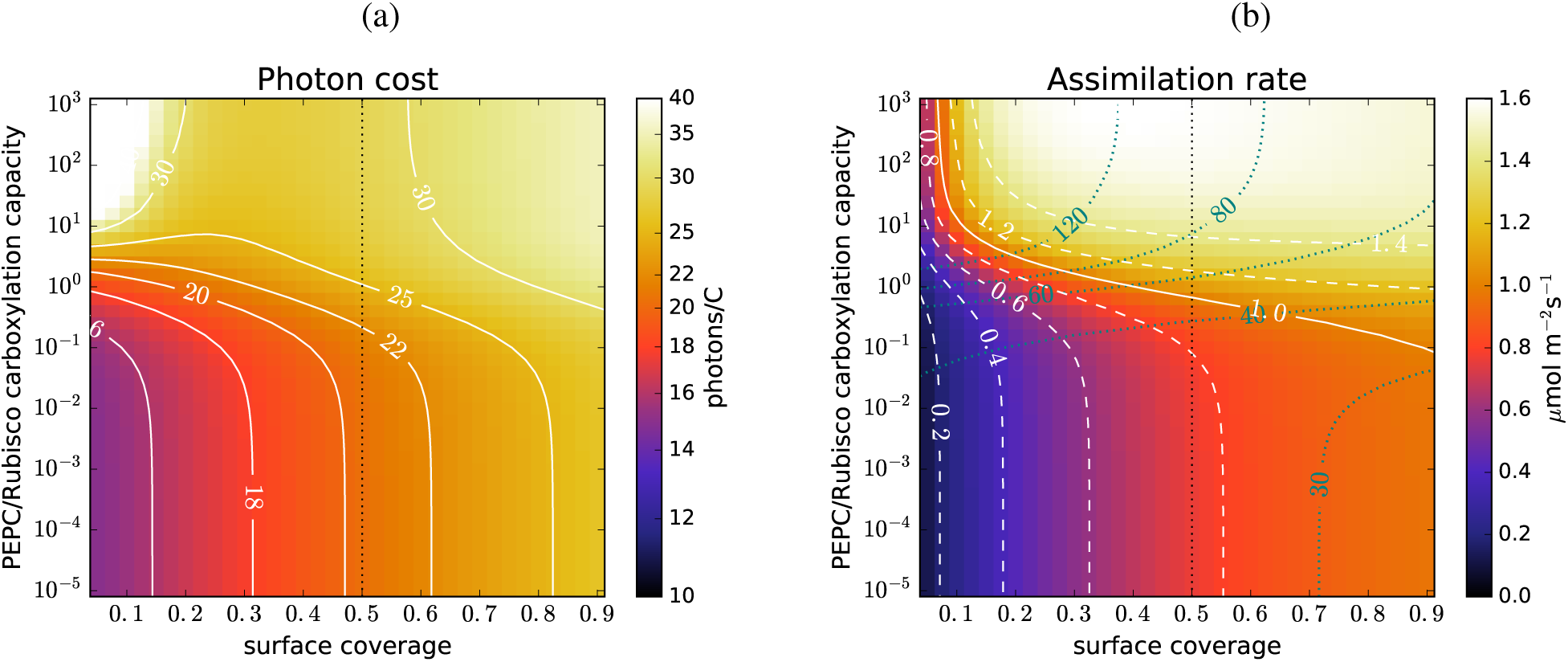
Plastid surface coverage and C_4_ photosynthesis: The photon cost (a) and the net assimilation rate (b) as functions of the chloroplast surface coverage and PEPC-to-RubisCO carboxylation capacity ratio, for the default parameter choice (Table 1). The purple lines in (b) mark the light-utilisation thresholds (in mM/s). The carboxylation capacity ratio is used instead of the PEPC concentration to quantify the C_4_ cycle activity because the cytoplasmic volume per plastid changes with the coverage. The vertical dotted line marks the surface coverage used as default in other figures.

Changing the permeability of the cell wall and plasmalemma (σ_*C*_) results in significant changes to the photon cost and the assimilation rate, as one could expect (Supplementary Figure 5). The efficacy of the C_4_ cycle (that is, its advantage or disadvantage over C_3_ photosynthesis) is only slightly affected, however. At very high σ_*C*_, the C_4_ cycle allows for a several-fold higher assimilation rate, as the bottleneck due to diffusion of CO_2_ through the cell wall is removed, but a concurrent increase in the photon cost means the plastid light-harvesting capacity would be limiting. This can be noted in the capacity thresholds which follow the assimilation rate isolines at high σ_*C*_ in Supplementary Figure 5(b).

**Figure 5:**
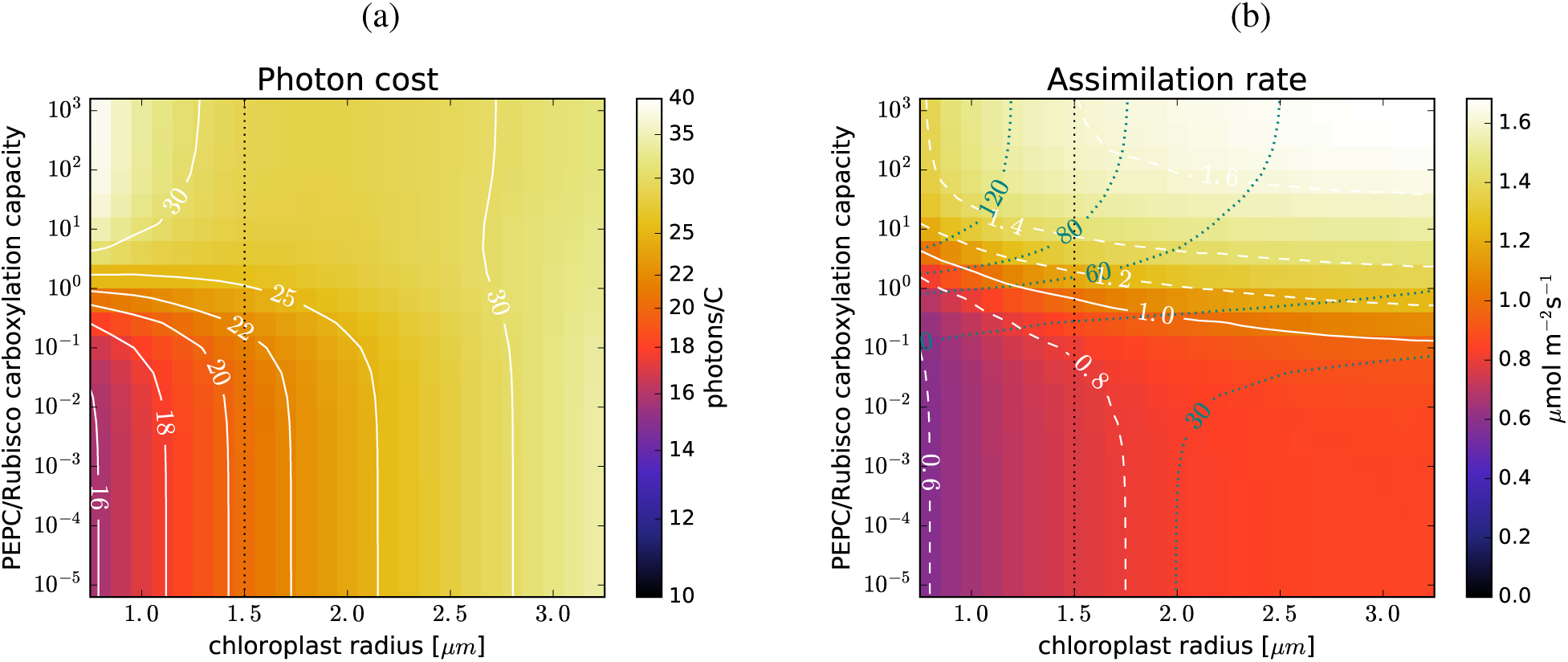
Plastid size and C_4_ photosynthesis: The photon cost (a) and the net assimilation rate (b) as functions of the chloroplast radius and PEPC- to-RubisCO carboxylation capacity ratio, for the default parameter choice (Table 1). The purple lines in (b) mark the light-utilisation thresholds (in mM/s). The carboxylation capacity ratio is used instead of the PEPC concentration to quantify the C_4_ cycle activity because the stromal volume per cell-surface area changes with plastid radius. The vertical dotted line marks the plastid size used as default in other figures.

Changing the chloroplast surface coverage (Figure 4) leads to an interesting result: while the activation of the C_4_ pump (when the envelope permeability is above the break-even threshold) will always cause the photon cost to rise,for lower surface coverages (30%-50%) this rise is less steep. A remarkable and non-intuitive consequence is that higher assimilation rate *per cell surface area* (and hence *per leaf-surface area*, assuming a fixed mesophyll-to-leaf surface ratio) can be achieved at lower chloroplast surface coverage, i.e. at a lower investment in plastids (Figure 4(b)).

Increasing the plastid size while keeping the cell surface coverage constant (Figure 5) means more RubisCO per cell surface area and hence a higher assimilation rate, but also a higher photon cost due to the increased RuBP oxygenation in the case of C_3_ photosynthesis. The C_4_ cycle, at high enough PEPC concentrations, can reverse this negative trend. At PEPC-to-RubisCO capacity ratios above 3, C_4_ efficiency increases with plastid size, although C_4_ photosynthesis still remains less efficient than C_3_ photosynthesis, except for unrealistically large plastids. This results in a higher assimilation rate per cell-surface area combined with lower demands on the light-harvesting capacity (Figure 5(b)).

### The gain under realistic light utilisation limits

We now examine what gains are achievable when the energy input is a constraining factor. This could be either due to limited light availability or limited light-harvesting capacity. We expect that at energy inputs below the level needed to operate C_3_ photosynthesis, activating the C_4_ pump would negatively affect the assimilation rate. Therefore we consider only situations where the energy constraints do not inhibit C_3_ photosynthesis. This will be the case at light-utilisation caps of 40 mM/s or more. If the thylakoid surface area is not the constraining factor in C_3_ photosynthesis, it should be possible to boost the plastid light-harvesting capacity well beyond 40 mM/s by overexpressing the photosystem complexes and associated proteins on the thylakoid. We examine photosynthesis under a realistic light- utilisation cap of ∼ 40 mM/s, and under a *reasonably* optimistic one of 80 mM/s. These correspond to a utilisation of 2% and 4% of the maximal incident solar flux on the plastid.^5^

Figure 6(a) shows how assimilation changes with the PEPC concentration at different envelope permeabilities, when the 40 mM/s cap is imposed. The steady-state operation is not affected as long as energy use remains below the cap. When energy availability becomes limiting there is a reduction in the concentrations of available RuBP- primed RubisCO and PEP-primed PEPC. As their carboxylation capacity ratio remains constant (by assumption), the operation under an energy constraint will be the same as unconstrained operation at a reduced, effective RubisCO concentration, with the same PEPC-to-RubisCO ratio. High PEPC concentration will then result in reduced assimilation, as the C_4_ cycle and Calvin-Benson cycle enzymes compete for energy resources. We might then expect that the optimal assimilation under an energy constraint is achieved exactly at the threshold where the energy usage reaches the cap. This is essentially the case for the 80 mM/s cap, but is not generally true. As the C_4_ cycle changes the operating conditions in the stroma (i.e. CO_2_ levels) this can result in a situation where a lower effective RubisCO concentration results in a higher net assimilation. The comparison of the assimilation gains (with respect to C_3_ photosynthesis) at the threshold PEPC concentration where the energy consumption reaches the cap, and at the concentration where the optimal assimilation is accomplished is shown in Figure 6(b). The respective PEPC concentrations are shown in Figure 6(c). It is evident that the C_4_ cycle activity has to be tuned to obtain the maximal benefit. Given that light supply fluctuates continually, dynamic control of the C_4_ cycle activity would have to be implemented. Alternatively, under-operating the cycle (i.e. having its activity level below the speculated optimum) may be a beneficial strategy.

**Figure 6:**
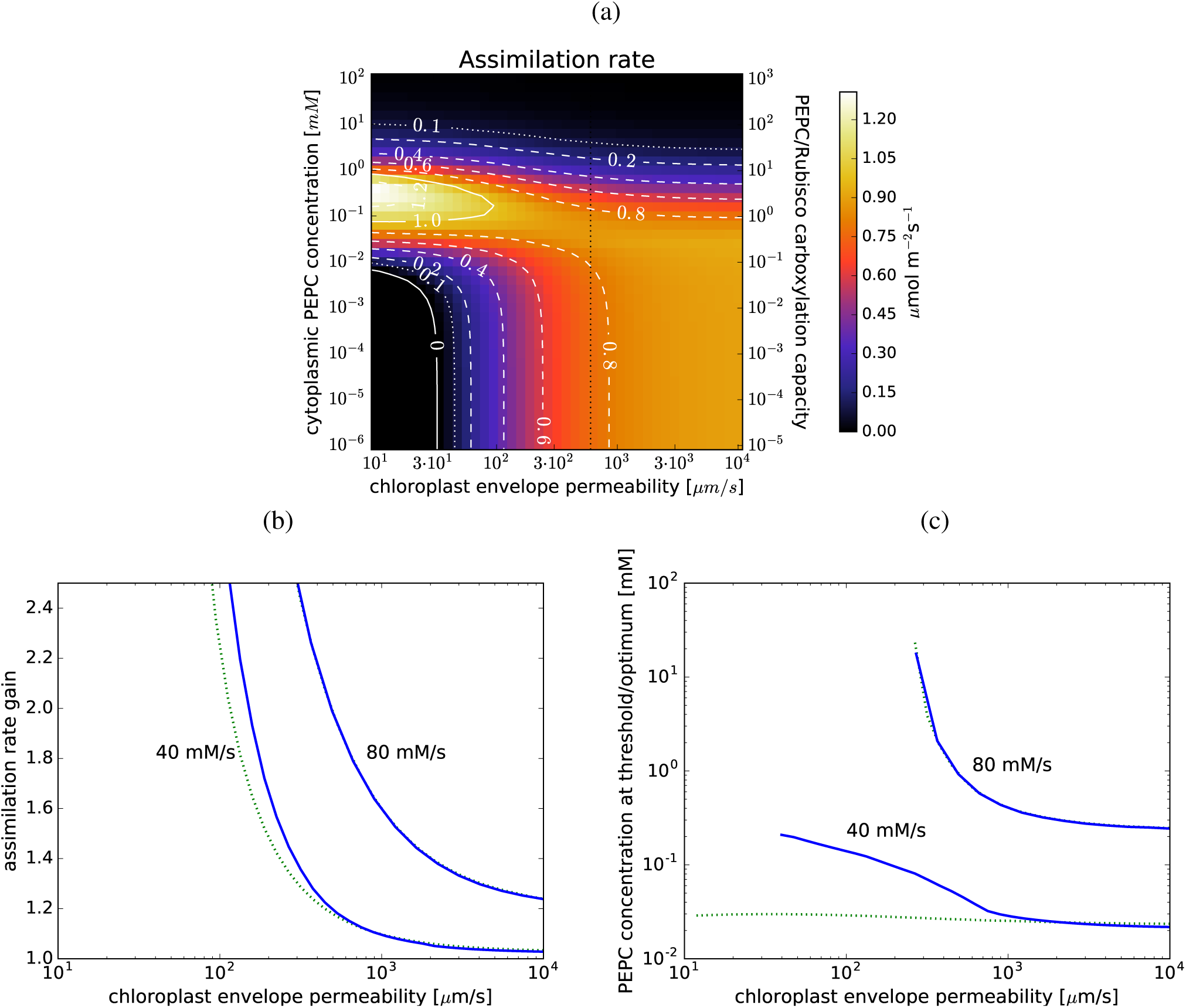
C_4_ photosynthesis at limited light-harvesting capacity. (a): The net assimilation rate as a function of the envelope permeability and PEPC concentration in the cytoplasm when the light input is capped at 40 mM/s. Parameters as in Fig. 3. The vertical dotted line marks the permeability used as default in other figures. (b): The relative gain in the assimilation rate at the PEPC activity levels where the light usage reaches 40 mM/s and 80 mM/s (dotted green lines, corresponding to the purple lines in Fig. 3(b)) and the maximal assimilation gains when the caps are imposed (blue lines). (c): the respective PEPC concentrations at which the optimal gains are achieved.

Even without a fine-tuned C_4_ cycle a sizeable gain in the assimilation rate can be expected as long as envelope permeability is not too large. Looking at the photosynthetic performance at the threshold where the energy consumption reaches the 40 mM/s cap, we predict that up to 20% gain in carbon assimilation at the permeability of 600 *µ*m/s may be achieved, with the photon cost rising by less than 10%. With the higher LHC of 80 mM/s (and sufficient sunlight) large gains are possible over the entire range of the envelope permeability values. Assimilation may even be doubled.

Stomatal conductance is continually tuned to the environment and when conductances are low photosynthesis is frequently CO_2_ deprived. Assimilation gains from using the C_4_ pump are much more prominent at low CO_2_ pressures in the internal airspaces, Figure 7(a). At 120 µbar CO_2_ the assimilation could be doubled, while still not exceeding the 40 mM/s light-utilisation cap (Figure 7(b)). In contrast, at 400 µbar no gain is possible with that energy cap (Figure 7(b)).

**Figure 7:**
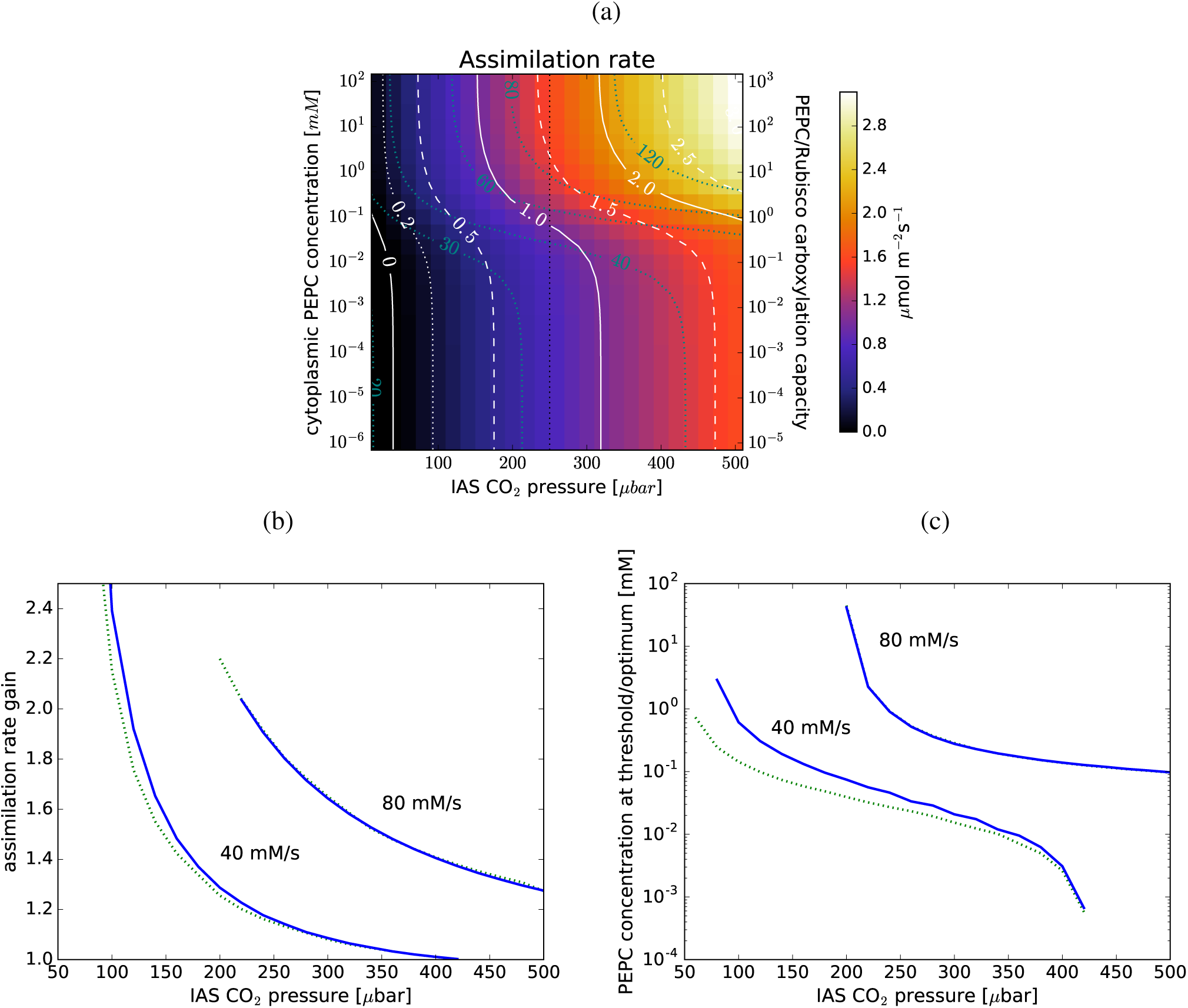
*C*_4_ photosynthesis at limited CO_2_ in the IAS. (a): The net assimilation rate as a function of the IAS CO_2_ pressure and PEPC concentration in the cytoplasm, for the default parameter choice (Table 1). No light utilisation cap is imposed, but the utilisation thresholds are marked in purple. The vertical dotted line marks the CO_2_ pressure used as default in other figures. (b): The relative gain in the assimilation rate at the PEPC activity levels where the light usage reaches 40 mM/s and 80 mM/s (dotted green lines) and the maximal assimilation gains when the caps are imposed (blue lines). (c): the respective PEPC concentrations at which the optimal gains are achieved.

## Discussion

We modelled a hypothetical cytoplasm-to-stroma C_4_ cycle in a C_3_ mesophyll cell geometry, and quantified carbon assimilation and photosynthetic efficiency. The proposed C_4_ pump would lead to an increase in assimilation rate whenever there is sufficient light-harvesting capacity *and* excess light is available. The magnitude of this gain is highly dependent on CO_2_ permeability of the chloroplast envelope and on operating conditions, such as the internal airspace CO_2_ pressure and light availability. At medium envelope permeability (600 *µ*m/s), CO_2_ pressure (250 µbar), and light-harvesting capacity (40 mM/s), the gain is moderate (20%). At low CO_2_ pressure (125 µbar), or at high light availability and harvesting capacity (80 mM/s), the gain becomes significant (85%), Figure 7(b). Due to the design of the model, which assumes optimal functioning of the C_3_/C_4_ enzymatic pathways, these predictions always represent the best case scenario. Even so, the massive predicted assimilation advantage under conditions of CO_2_ deprivation is unlikely to be spurious. As CO_2_ deprivation is a common hazard facing plants in dry and warm climates - which are typically well-lit - the development of the proposed C_4_ pathways could be *very* beneficial for creating drought-resistant high-yield crop strains. Regarding this it is interesting to note that terrestrial species that *have* evolved single-celled C_4_ photosynthesis grow in salty depressions in semi-arid regions - the conditions that would likely lead to low CO_2_ within the leaf (Voznesenskaya et al., 2001, 2002).

Ideally a regulation mechanism should be incorporated to moderate the activity of the C_4_ pump based on the energy availability, so as to prevent it from competing adversely with the Calvin-Benson cycle. Regulation of the C_4_ cycle based on the ambient light levels and CO_2_ availability is already present in Kranz-type C_4_ plants (Furbank and Taylor, 1995), so simply implementing existing C_4_ regulatory mechanisms may allow this.

Our conclusions are generally in qualitative agreement with von Caemmerer (2003), but our detailed examination of parameter space produces more positive results, inspiring greater optimism. As Figure 6(b) shows, predicted gains from the C_4_ cycle are much higher at lower envelope permeabilities. Likewise, although we agree with von Caemmerer (2003) that the C_4_ cycle will be cost-inefficient, the difference between carbon assimilation costs in C_3_ and C_4_ photosynthesis is smaller at lower envelope permeability, and, as we demonstrate by evaluating the expected assimilation at limited light-harvesting ability, the operation of a C_4_ cycle need not be prohibitively expensive.

To elucidate the differences in our conclusions, we attempt a more direct comparison with the results of von Caemmerer and Furbank (2003). At 200 ppm CO_2_ in the IAS, they predict that operating the C_4_ pump at 1:1 PEPC- to-RubisCO carboxylation capacity ratio would result in a 40% increase in the assimilation rate and a 70% increase in energy cost per assimilated carbon (Figure 5 in von Caemmerer and Furbank (2003)). Their model expresses gas conductances and enzyme catalytic capacities per leaf-surface area, so a comparison requires an assumption of the mesophyll-to-leaf surface area ratio. Using a ratio of 13.5 (the value is similar to the ratios found for A. thaliana (8-10) by Tholen et al. (2008)), the RubisCO catalytic capacities in the two models match,^6^ so we use this value for the comparison. Their conductances then correspond to permeabilities of the envelope, and of the cell wall and plasmalemma, of approximately 10^3^ *µ*m/s each. With the same parameters we get a 50% increase in the assimilation rate with a 30% increase in the photon cost (from 17/C to 22/C). There is a significant difference in the predictions of the energy cost of C_4_ photosynthesis. The difference likely stems from different accounting methods - von Caemmerer and Furbank (2003) consider ATP consumption whereas our quantification in terms of light-use factors in the fact that the C_4_ cycle does not need a reductive agent so its ATP requirements can be met by the cyclic electron transfer chain, which is more efficient at creating a proton gradient across thylakoid membrane, and thus ATP production per photon is higher.

Another promising result is that the pathway’s beneficial effects can be increased further by reducing the chloroplast surface coverage, bringing it into the region in Figure 4(a) where the rise in the photon cost when the C_4_ pump is active is less pronounced. This minor change to the cell anatomy would allow for the same assimilation rate to be achieved with a reduced plastid investment, translating into an even higher plant growth rate. One way this could be accomplished might be to arrest or slow down the chloroplast division cycle. A possible side-effect would be an increase in the average plastid size, which would further benefit C_4_ photosynthesis (Figure 5). An illustration of possible benefits from a design strategy that combines the implementation of a C_4_ cycle with alterations in the chloroplast surface coverage is presented in Figure 8. The design steps are broadly outlined in Figure 8(b). Figure 8(a) shows how the assimilation rate varies with the surface coverage (assuming no changes in the plastid size) for C_3_ photosynthesis, and C_4_ photosynthesis at 40 mM/s and 80 mM/s light utilisation thresholds (compare with Figure 4(b)). Starting with C_3_ photosynthesising plastids at 50% cell surface coverage (a_0_), implementing the C_4_ pump and boosting the light-harvesting capacity to 40 mM/s (a_1_) or 80 mM/s (a_2_) would result in a 15% or an 85% increase in the assimilation rate respectively. Alternatively, at 40 mM/s light-harvesting capacity, the number of plastids could be reduced by 20% (b_1_) without any loss in assimilation compared to C_3_ photosynthesis. Boosting the light-harvesting capacity to 80 mM/s would allow for a 65% increase in the assimilation rate with 20% fewer plastids (b_2_), or alternatively, for a 60% reduction in the number of plastids without a decrease in assimilation (c_2_). If the plastids are also enlarged in the process, even larger gains may be possible. The level of required C_4_ cycle expression, quantified by the PEPC/RubisCO carboxylation capacity ratio, would not need the exceed the observed level of C_4_ cycle activity in C_4_ plants (2-7; von Caemmerer et al. (2014)), even at 80 mM/s light-harvesting capacity (Figure 8(a)). The relative expression of the two photosystems may need to be rebalanced however, to allow for a larger cyclic electron current through PS-I (Figure 8(a)). The optimal modification strategy would be the one that maximises the return on resource investment. To calculate this however, the maintenance costs also need to be established. Quantifying the return-on-investment and deciding the optimal strategy will require additional research

**Figure 8:**
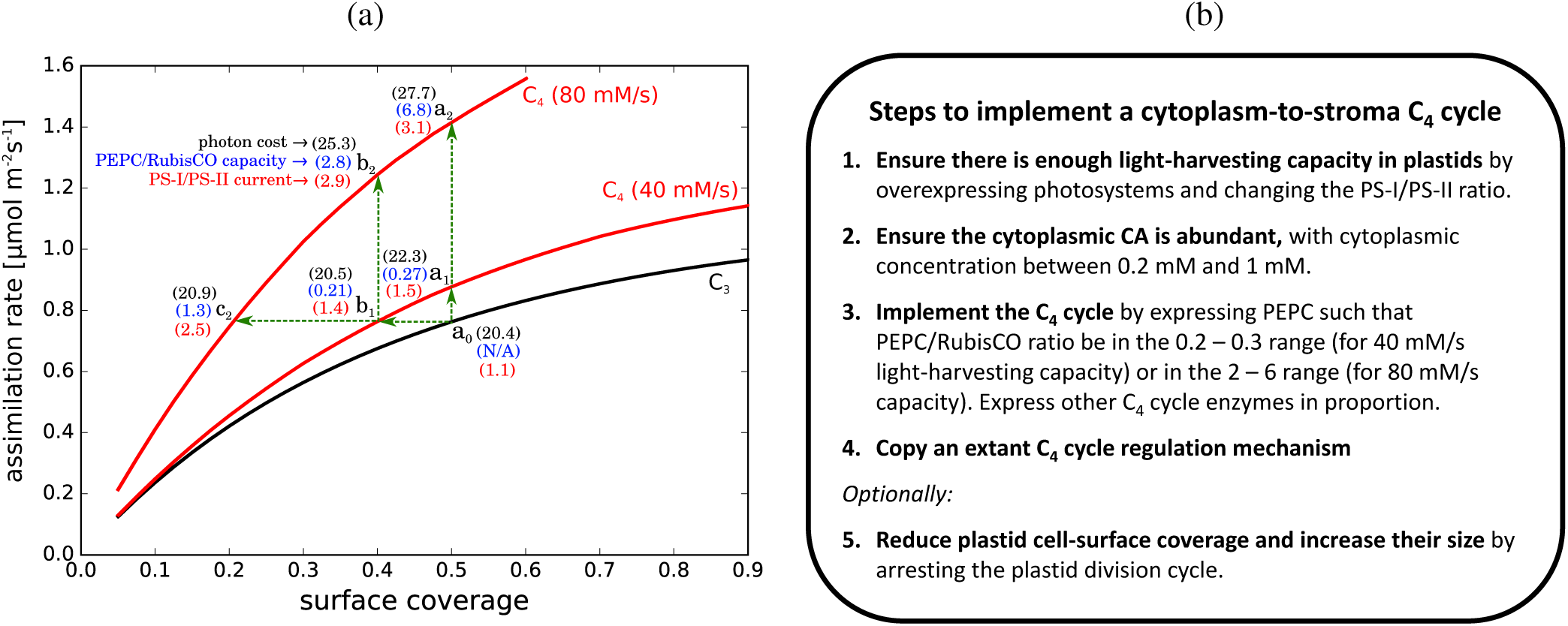
Altering plastid surface coverage and light harvesting capacity: (a) The assimilation rate per cell surface area as a function of chloroplast surface coverage in the case of C_3_ photosynthesis (black line) and C_4_ photosynthesis at the C_4_ cycle activity levels where light-use reaches 40 mM/s and 80 mM/s thresholds (red lines). The numbers in parentheses show the respective photon costs (black), PEPC-vs-RubisCO carboxylation capacity ratios (blue), and ratios of electron current through PS-I and PS-II (red). Green arrows illustrate organism modification strategies discussed in the main text. Parameters as in Fig. 4. (b) An outline of a recipe for making a functional C_4_ photosynthesising prototype.

## Acknowledgements

This research was supported by the BBSRC project grants BB/M011291/1, BB/I024445/1, BB/M011356/1, and BB/M01133X/1.

## Supplementary Information

### Mathematical outline of the model

To find the gas currents under steady-state photosynthesis we need to solve the system of stationary diffusion-reaction equations for position-dependent concentrations of oxygen, carbon-dioxide, and bicarbonate - *n*_*O*_, *n*_*C*_, and *n*_*biC*_ - satisfying appropriate boundary constraints and flux-balance conditions. The equations are of the form

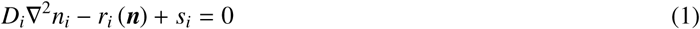

where the index *i* stand for *C, O*, and *biC* respectively. *D*_*i*_ is the compartment-dependent diffusion coefficient. *r*_*i*_ and *s*_*i*_ are the reaction and source terms. Both will depend on the location (the compartments, i.e. stroma, cytoplasm, and vacuole, or the intramembrane and intraenvelope spaces). The reaction term may in principle depend on the local value of any of the three concentrations, which we subsume into a ‘vector’ form ***n*** ≡ (*n*_*O*_, *n*_*C*_, *n*_*biC*_). The source terms are determined by flux-balance conditions to be addressed later. As all of the terms depend on the compartmental location, we introduce characteristic functions χ_*P*_(*r*), χ_*C*_(*r*), and χ_*V*_ (*r*), which are equal to one if *r* is respectively within the plastid, cytoplasm, or vacuole, and zero otherwise. This way, we can specify the reaction terms as

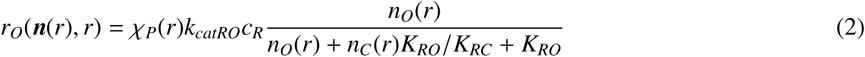

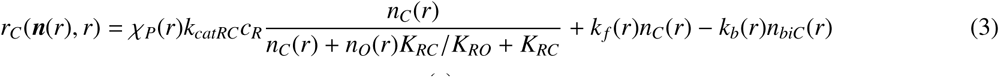

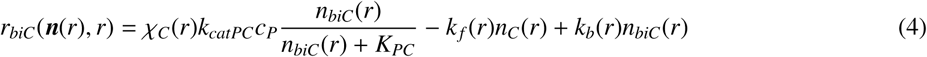

where we used stationary Michaelis-Menten forms for the competitive reaction of RuBP-primed RubisCO with O_2_ and CO_2_, and for the reaction of the bicarbonate with PEPC. *k* _*f*_ and *k*_*b*_ are the forward and backward reaction rates for the CO_2_ to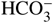 interconversion. They will depend on the local pH value and on the presence or absence of the anhydrase. They are given by Johnson (1982) as^1^

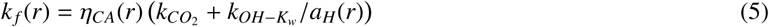

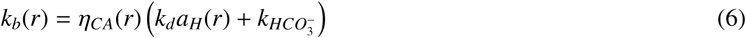

*a*_*H*_(*r*) is the proton activity, given by the local pH, *a*_*H*_ = 10^-*pH*^ M, and *η*_*CA*_ is the CA-dependent reaction boost factor. It is equal one where CA is absent (e.g. vacuole), and to a large number (10^6^ by default) where CA is present (i.e. in the stroma and cytoplasm). We do not allow for the CO_2_-bicarbonate interconversion in the intramembrane and intraenvelope space (since it is an hydrophobic environment), so *k* _*f*_ and *k*_*b*_ are set to zero there.

The source terms *s*_*i*_ reflect the release of O_2_ and CO_2_ as products of the relevant chemical pathways connected to photosynthesis. They are determined by the input the same gasses as reactants in those pathways. We first define the cumulative fluxes.

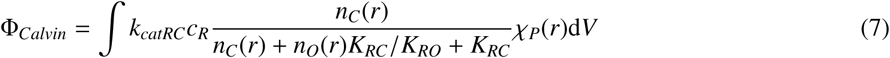

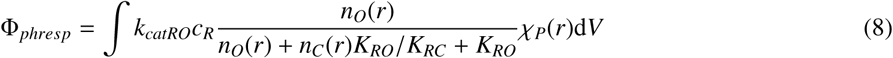

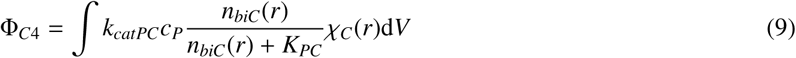

The source terms are

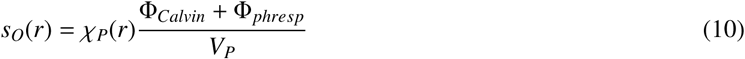

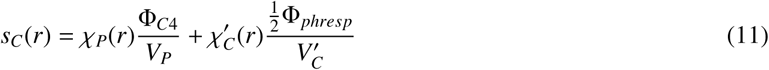

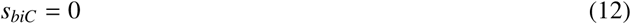

where *V*_*i*_ stand for the volumes of particular compartments,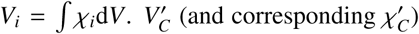 stands for the part of the peripheral cytoplasmic space accessible to the mitochondria.

The flux-balance relations incorporated in the source terms address the Hill reaction at the thylakoid (one oxygen molecule created for each CO_2_ and O_2_ molecule reacting with RuBP matches the NADPH creation through linear electron transfer chain with its consumption in the Calvin cycle and photorespiration), the photorespiratory CO_2_ release in the mitochondria (one CO_2_ molecule released for every two RuBP oxygenation events), and the release of CO_2_ from the C_4_ acid decarboxylation in the stroma (one CO_2_ molecule for each 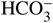 ion reacting with PEP in the cytoplasm).

The diffusion constant of a specie in a particular compartment, *D*_*i*_, is equal to the diffusion constant of that specie in water *D*_*i,aq*_ divided by the viscosity of the liquid filling the compartment

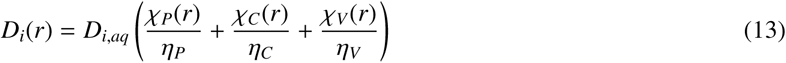

However, within the tonoplast membrane and the chloroplast envelope, the diffusion coefficient is set to reflect the permeability of the particular barrier. For a barrier with thickness *θ* and permeability σ_*i*_, we’ll have

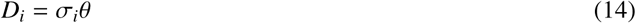

Diffusion through the cell wall and plasmalemma is modelled by a boundary condition connecting the current density of CO_2_ and O_2_ perpendicular to the boundary surface with the difference between the local and equilibrium gas concentrations:

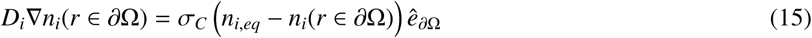

Here *i* stands for oxygen or carbon-dioxide (we assume the bicarbonate cannot cross the plasmalemma), ∂Ω is the cell boundary surface and 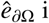 is the unit vector perpendicular to that surface. σ_*C*_ is the combined permeability of the cell wall and the plasmalemma, while *n*_*i,eq*_ is the stationary dissolved concentration of carbon-dioxide/oxygen in the thin wetting layer outside the cell wall, which is presumed to be in thermal equilibrium with the pressure of the respective gas in the internal airspace.

At other boundary surfaces we assume von Neumann boundary conditions, i.e. there is no current in or out of the simulated region

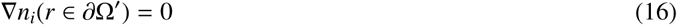

We have posed the model in a very general way as a system of nonlinear partial integro-differential equations in three dimensions. In reality we seize the advantage of the postulated cylindrical symmetry of the system. The resulting problem, which is effectively two-dimensional, is solved iteratively by a finite element method on a prespecified simplex mesh. The mesh is algorithmically constructed to follow the natural boundaries of the simulated system (i.e. the internal and external surface of the envelope and the tonoplast membrane). We use DUNE/PDELab libraries with BCGS solver on P_2_ elements (Blatt et al., 2016; Alkämper et al., 2016). As nonlinear PDE’s require iterative solving, there is a natural way to include our integrative flux-balance conditions by updating the source terms with each iteration.

The light-limited operation can be simulated by also evaluating the energy usage *W*^*n*^ during each iteration and scaling the concentration of RubisCO and PEPC, if the usage exceeds some capping threshold *W*_*cap*_.

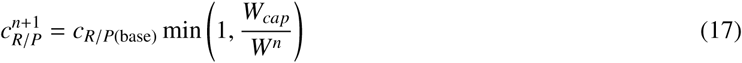

where we denoted the iteration number in the superscript. The light usage is evaluated from the overall fluxes in the energy consuming biochemical pathways.

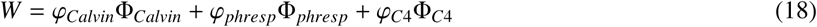

where *ϕ*_*i*_ stand for the photon cost of RuBP and PEP regeneration after each carboxylation or oxygenation event; and Φ_*i*_ are the respective fluxes.

The net carbon assimilation rate is determined by the competition of the Calvin and photorespiratory pathways:

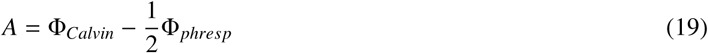

and the net photon cost of carbon assimilation (i.e. the inverse quantum efficiency of the photosynthesis) is

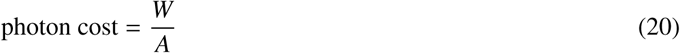

The assimilation shown in the figures is the assimilation rate per cell surface. It is obtained by dividing *A* with the base of the simulated cylinder

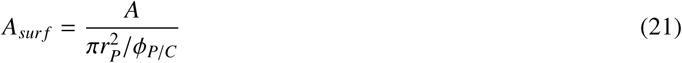

where *Φ*_*S* *C*_ is the chloroplast surface coverage.

## Supplementary Figures

**Figure 1:**
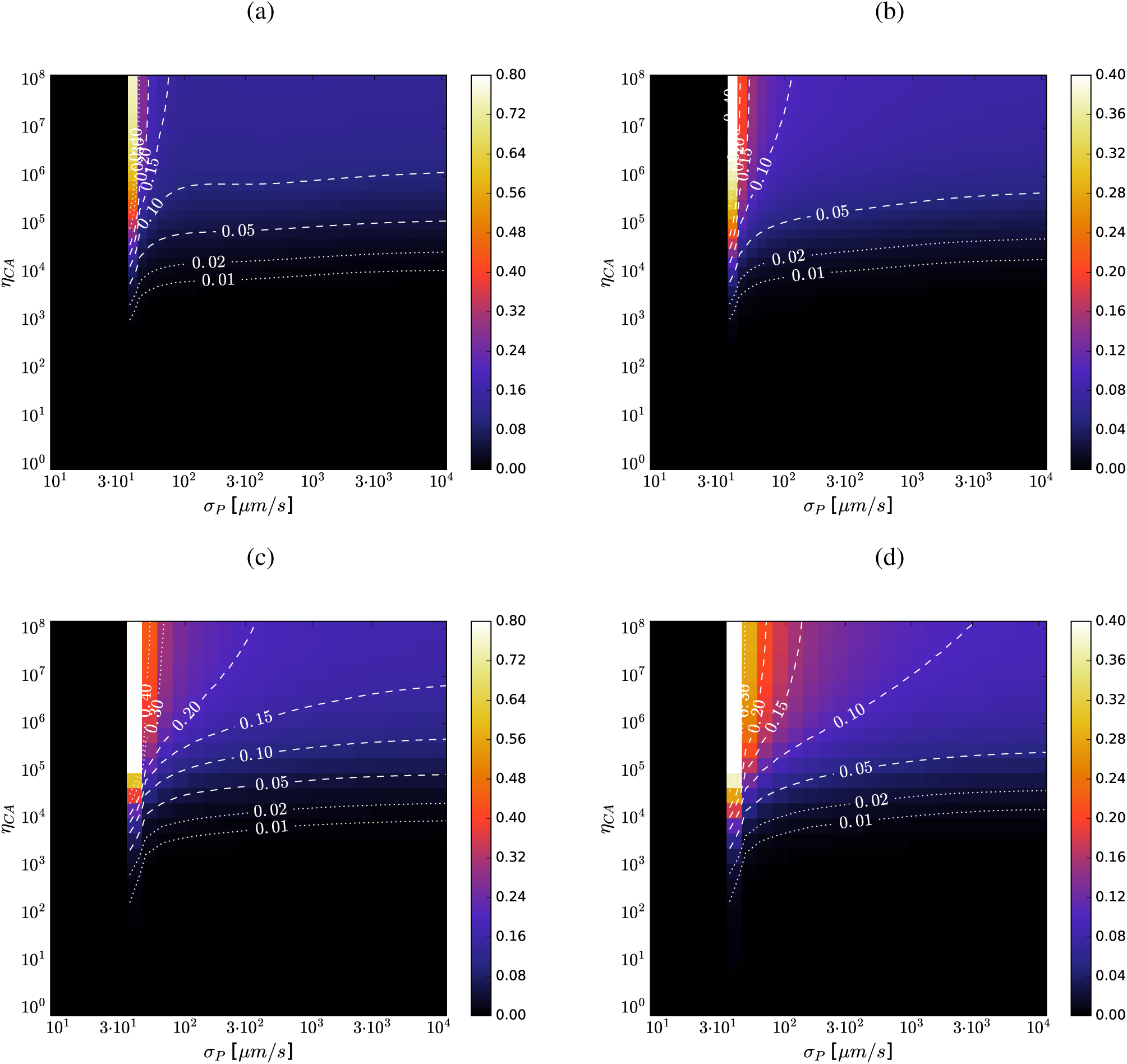
CA efficacy vs. envelope permeability in C_3_ photosynthesis. (a) and (b): Relative increase in carbon assimilation rate and reduction in photon cost due to presence of CA in the chloroplast stroma, as functions of the envelope permeability (*σ*_*P*_) and the 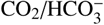interconversion rate boost (*η*_*CA*_); for *p*_*CO*_2 = 250 *µ*bar and *σ*_*C*_ = 200 *µ*m/s. (c) and (d): Same as (a) and (b) for the case when CA is present both in the stroma and in the cytoplasm.

**Figure 2:**
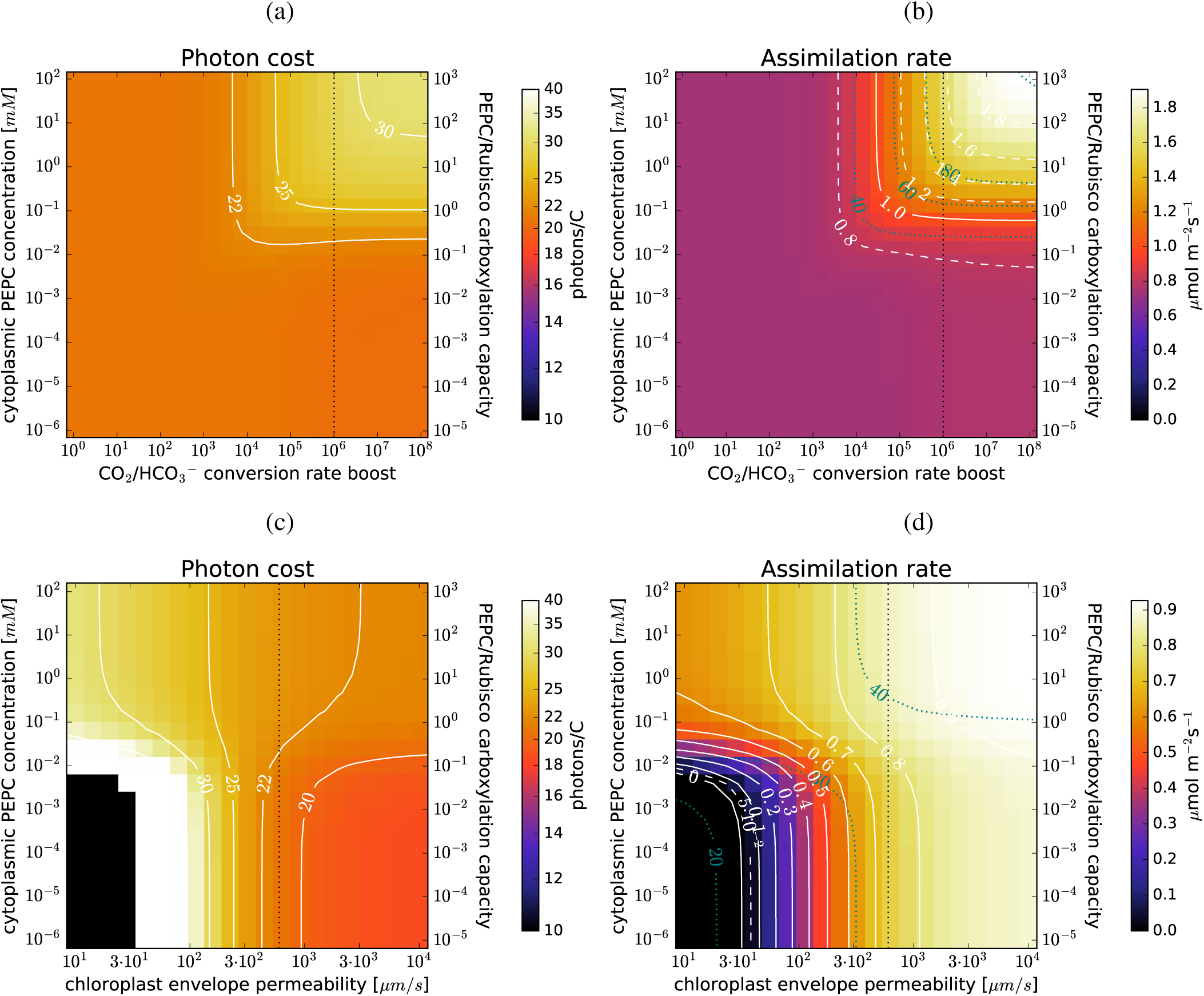
CA efficacy and C_4_ photosynthesis: (a) and (b): the photon cost and the net assimilation rate as functions of the 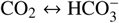 conversion rate boost due to cytoplasmic CA (*η*_*CA*_) and PEPC concentration in the cytoplasm (*c*_*P*_), for the default parameter choice (Table 1). (c) and (d): the photon cost and the net assimilation rate as functions of the envelope permeability (*σ*_*P*_) and PEPC concentration in the cytoplasm (*c*_*P*_), when cytoplasmic CA is insufficient (*η*_*CA*_ = 10^4^). The vertical dotted lines mark the values used as defaults in other figures.

**Figure 3:**
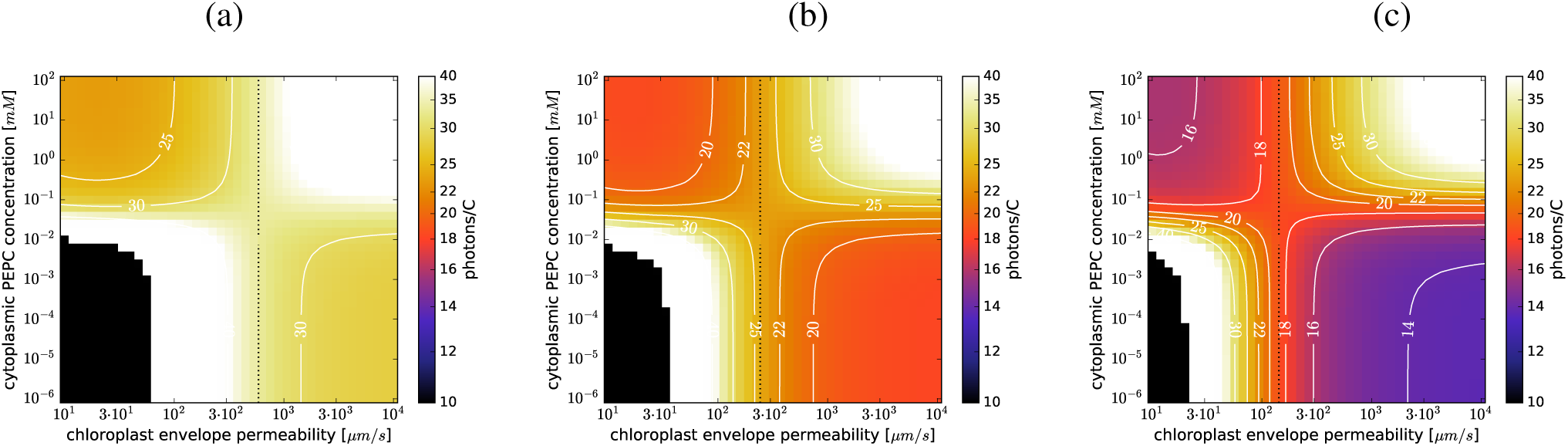
*C*_4_ photosynthesis at different IAS CO_2_ levels: The photon cost as a function of the envelope permeability (*σ*_*P*_) and PEPC concentration in the cytoplasm (*c*_*P*_) at *p*_*CO*_2 = 150 *µ*bar (a), *p*_*CO*_2 = 250 *µ*bar (b), and *p*_*CO*_2 = 400 *µ*bar (c). The dotted vertical marks the threshold envelope permeability below which the C_4_ cycle is cost-efficient.

**Figure 4:**
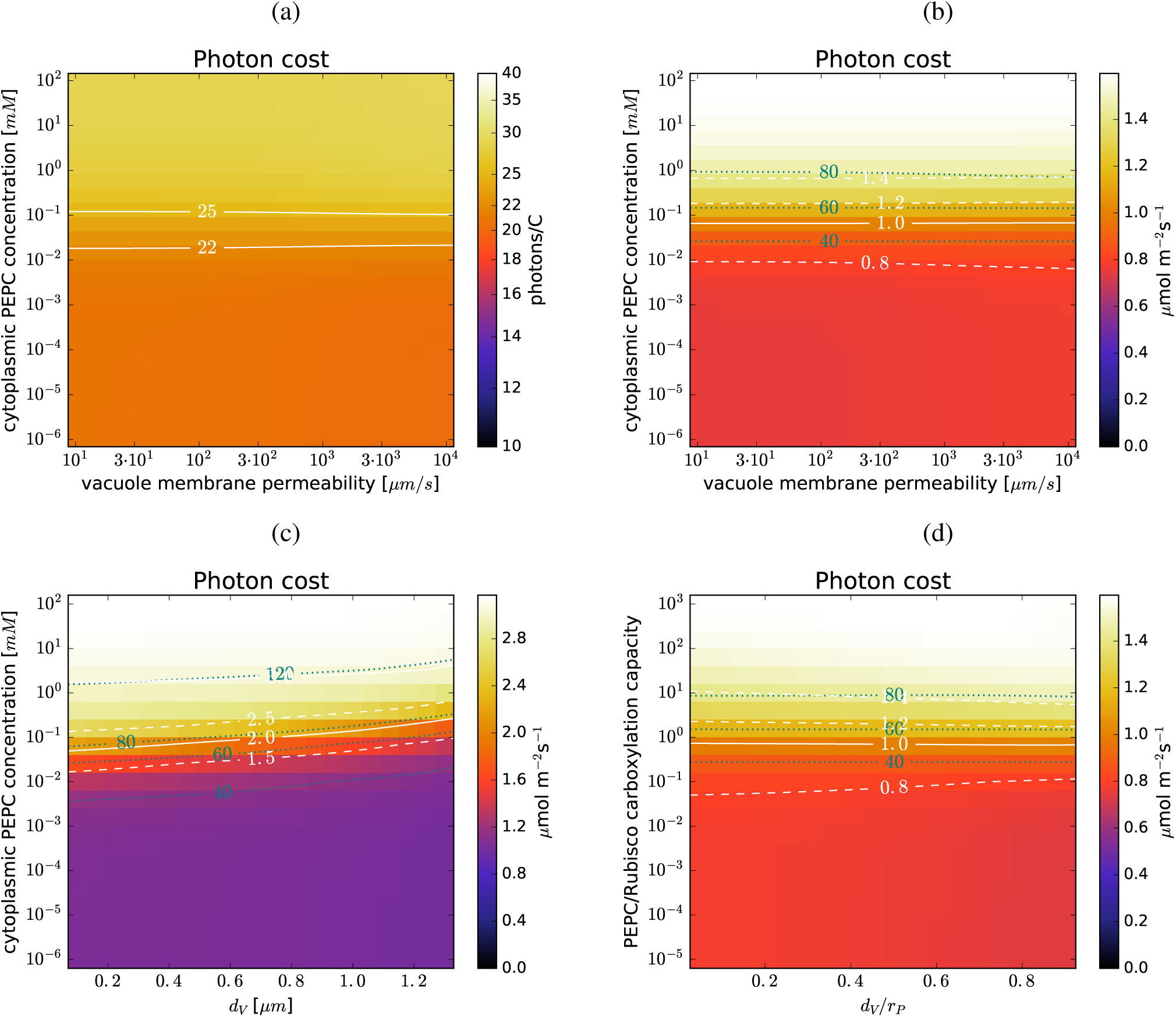
Vacuole impact on C_4_ photosynthesis: (a) and (b): The photon cost and the assimilation rate as functions of the vacuole membrane permeability *σ*_*V*_ and the cytoplasmic PEPC level (*c*_*P*_), at *σ*_*C*_ = 200 *µ*m/s, *σ*_*P*_ = 600 *µ*m/s, and *p*_*CO*_2 = 250 *µ*bar. (c) and (d): The assimilation rate as a function of the drop of the vacuole (*d*_*V*_) (which reduces the cytoplasmic volume) and the PEPC concentration (c) or the PEPC-to-RubisCO carboxylation capacity ratio (d). Parameters same as above, with *σ*_*V*_ = 2*σ*_*P*_ = 1200 *µ*m/s.

**Figure 5:**
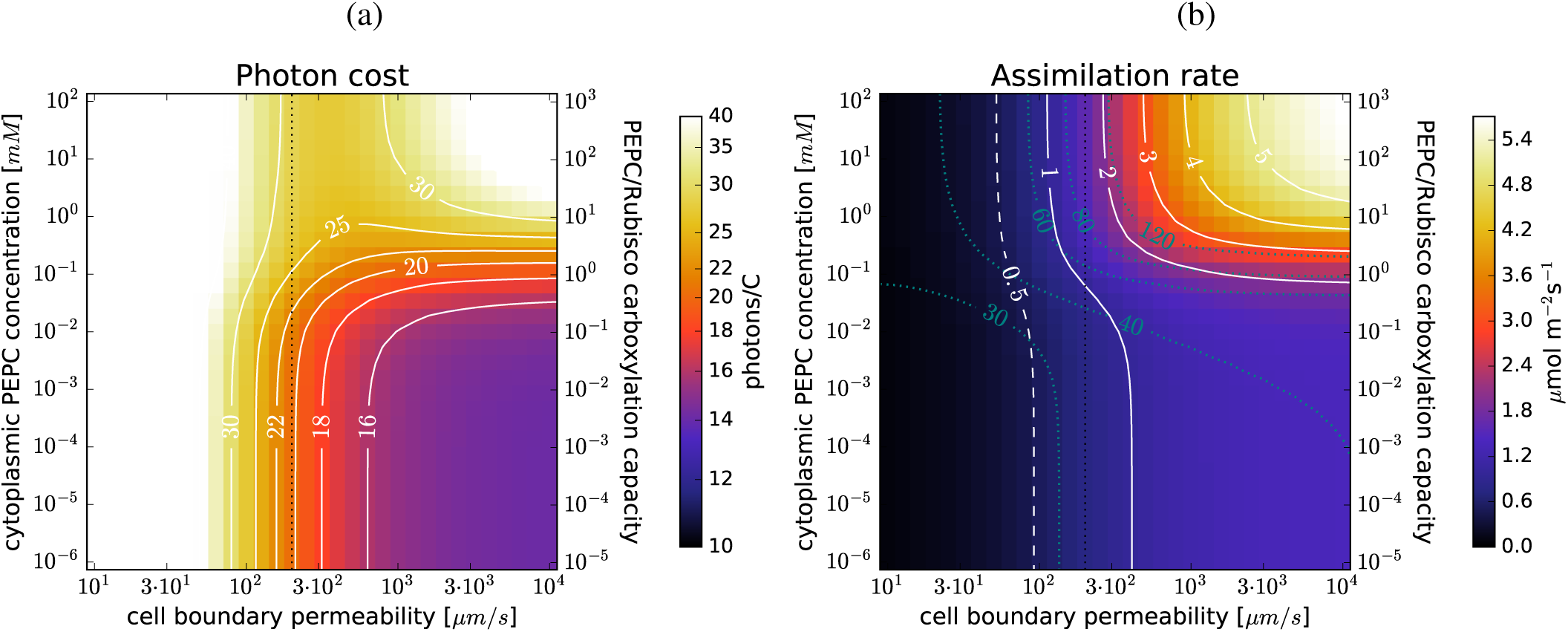
Cell wall permeability and C_4_ photosynthesis: (a) and (b): the photon cost and the net assimilation rate as functions of the combined permeability of the cell wall and plsmalemma (*σ*_*C*_) and PEPC concentration in the cytoplasm (*c*_*P*_), for the default parameter choice (Table 1). The vertical dotted line marks the permeability used as default in other figures.

## Footnotes

1 At 50% plastid surface coverage, this corresponds to 0.5 % of maximal solar flux passing through the cell surface.

2 The effective RubisCO and PEPC concentrations are scaled proportionally, so their carboxylating capacity ratio stays fixed. A plant with optimised control mechanisms could in principle independently alter the activity of the two carboxylases under light deprivation, but we do not assume the presence of such control mechanisms.

3 Here we only vary the efficacy of cytoplasmic CA, while keeping the efficacy of stromal CA fixed at 10^6^.

4 Depending on the particular values of *η*_*CA*_ and the permeability of the cellular boundary (σ_*C*_), one or the other will form a bottleneck first. The relevant rates are the 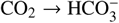 conversion rate and the volume-adjusted rate of CO_2_ diffusion from IAS (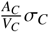,where *A*_*C*_ is the cell surface area and *V*_*C*_ is the volume of the peripheral cytoplasm). For default choice of parameter values (including *η*_*CA*_ = 10^6^), these are roughly 4 10^4^ s^−1^ and 500 s^−1^, so the diffusion of CO_2_ from the internal airspaces is limiting. At *η*_*CA*_ = 10^4^ the conversion rate is only 400 s^−1^ so it becomes limiting instead.

5 Or 1% and 2% of solar flux incident on cell surface (at 50% chloroplast surface coverage).

6 The RubisCO activity in von Caemmerer and Furbank (2003) is 100 *µ*mol/m^2^s, and the gas conductances of the envelope and the cell wall are 0.8 mol/bar m^2^s each.

1 The values of the individual rates are *k*_*CO*2_ = 0.037 s^−1^, *k*_*OH*-*Kw*_ = 7.1 · 10^−11^ Ms^−1^, *k*_*d*_ = 7.6 · 10^4^ M^−1^s^−1^, and *k*_*HCO*_- = 1.8 · 10^−4^ s^−1^ (Johnson, 1982).

